# Estimating the Lambda measure in multiple-merger coalescents

**DOI:** 10.1101/2023.03.10.532088

**Authors:** Verónica Miró Pina, Émilien Joly, Arno Siri-Jégousse

## Abstract

Multiple-merger coalescents, also known as Λ-coalescents, have been used to describe the genealogy of populations that have a skewed offspring distribution or that undergo strong selection. Inferring the characteristic measure Λ, which describes the rates of the multiple-merger events, is key to understand these processes. So far, most inference methods only work for some particular families of Λ-coalescents that are described by only one parameter, but not for more general models. This article is devoted to the construction of a non-parametric estimator of the density of Λ that is based on the observation at a single time of the so-called Site Frequency Spectrum (SFS), which describes the allelic frequencies in a present population sample. First, we produce estimates of the multiple-merger rates by solving a linear system, whose coefficients are obtained by appropriately subsampling the SFS. Then, we use a technique that aggregates the information extracted from the previous step through a kernel type of re-construction to give a non-parametric estimation of the measure Λ. We give a consistency result of this estimator under mild conditions on the behavior of Λ around 0. We also show some numerical examples of how our method performs.

## 1 Introduction

Inferring the evolutionary history of a sample of a present-time population is a very challenging task for both biological and mathematical communities. If no information is available on the past of the population, it is natural to consider Markov processes to model its genealogical tree. The family of Λ-coalescents, also called multiple-merger coalescents [Donnelly and Kurtz, 1999, Pitman, 1999, Sagitov, 1999], provides a rich class of processes to this purpose. They appear naturally as limit genealogies in population models, with possibly highly variable reproductive success [Sagitov, 1999]. Their flexibility (and richness) comes from the fact that their dynamics are fully described by a measure Λ on the interval [0, 1], in the sense that Λ determines all the rates of multiple-merger coalescences. The intrinsic exchangeability of the branches at a given time in Λ-coalescents makes them particularly suited for modeling genealogies in a very general framework. The family of Λ-coalescents contains the Kingman coalescent [Kingman, 1982b,a], which is considered as the neutral model for random genealogies. We refer to Berestycki [2009], Gnedin et al. [2014] for a mathematical survey that contains most of the justifications of the classical formulas that we use in this paper.

Some subclasses of Λ-coalescent, such as Beta-coalescents [Schweinsberg, 2003] are intensively used in practice since they are quite easily interpretable and are characterized by only one or two parameters. The inference task is then reduced to the estimation of this parameter, that represents the level of reproductive skewness in the population. This family is now well admitted to fit to populations with highly skewed offspring distribution, e.g. marine species [Sigurgíslason and Á rnason, 2003, Niwa et al., 2016], tuberculosis cells [Menardo et al., 2020] or cancer cells [Kato et al., 2017]. Additionally, the class of Beta-coalescents contains the Bolthausen-Sznitman coalescent that provides a genealogical process for modeling populations under strong selection effects [Neher and Hallatschek, 2013, Desai et al., 2013, Schweinsberg, 2017, Cortines and Mallein, 2017, Schertzer and Wences, 2023]. Another class of coalescents that has been used for modelling populations with skewed offspring distribution are the Dirac-coalescents [Eldon and Wakeley, 2006], which are characterized by two parameters.

Most of the previous inference work on Λ-coalescents has focused on estimating the characterizing parameters of the one parameter families cited above [Birkner et al., 2013, Eldon et al., 2015, Koskela, 2018, Hobolth et al., 2019]. But the inference of more general Λ measures is still a challenging issue. To our knowledge, it was only treated in Koskela et al. [2018], with Bayesian non-parametric methods. The authors consider a forward in time model of type frequencies, more adapted to data sampled at multiple time points, that are difficult to obtain from natural populations. In this paper we present an estimator that uses only contemporaneous observations.

Note that the family of Λ-coalescents does not contain more general genealogical models for populations with varying size [Freund, 2020], under bottlenecks [González Casanova et al., 2021], with diploid reproduction mechanism [Birkner et al., 2018], or recombination, which are part of a larger class of coalescents, offering more possibilities for inference and model selection (see for example [Eldon et al., 2015, Blath et al., 2016, Koskela, 2018, Sainudiin and Véber, 2018, Koskela and Wilke Berenguer, 2019, Freund and Siri-Jégousse, 2021].

In this work, we use approximations via Bernstein polynomials to establish a non-parametric estimator of the density of Λ, which is built from estimates of the multiple merger rates of the Λ-coalescent process. These estimates are the solutions of a linear system where the coefficients are the expected branch lengths (or SFS) estimated from different resamplings of the observed genomic data of the sample. This technique relies on generalizations of classical recursions for functionals of the coalescent (see Proposition 2.1). By reconstructing the density function of the measure Λ, and by visualizing it, our technique should help to detect some unexpected behavior of the population and to reject a whole class of coalescent models, e.g. Beta coalescents.

The type of observations that we use are SNP^1^ matrices that summarize mutation data: at every locus (column) where a mutation is observed, the individuals (lines) are assigned a 1 if they carry the mutation and a 0 otherwise (see Table 2 as an example). This means that the method can only work for polarized data i.e. data where derived and ancestral states can be tell apart. We assume that mutations at each locus occur independently. Notice that this does not mean that the columns of the SNP matrix are independent of each other, since the observed mutations all depend on the evolutionary history of the sampled population. From the SNP matrix we can compute the SFS, which is the vector such that its *i*-th component contains the number of observed mutations shared by exactly *i* individuals in the sample. This is a statistic widely used in population genetics, both for model selection [Eldon et al., 2015, Freund and Siri-Jégousse, 2021] and for parameter inference [Fu, 1995, Kersting et al., 2021]. In this paper, we use an extension of the SFS, the so-called *weighted SFS*, which can also be read from the SNP matrix by subsampling the individuals (lines), as explained in section 2.3. In the case of diploid (or polyploid) species, our method requires phased data so that each line of the SNP matrix would represent a haplotype. In addition, we assume given a collection of SNP matrices *S*_1_, …, *S*_*n*_ governed by the same subjacent Λ-coalescent. We also assume that those matrices are independent (in the probabilistic sense) so that no coalescence in one genealogical branch of one matrix *S*_*i*_ have an influence on the other matrices. For example, one can think of SNP matrices obtained from different replicates in an evolutionary experiment or from multiple isolated populations, that have evolved independently, under similar conditions (see for example Lenski et al. [1991] for *E. coli*, Johnson et al. [2021] for yeast or Teotónio et al. [2017] for *C. elegans*). For applications to real populations, it is not possible to obtain independent SNP matrices from the same population. However, ignoring dependencies between genomic regions that are at sufficient recombination distance is a natural approximation and it has been used for example in McVean and Cardin [2005] to define the sequentially Markov coalescent. In the case of multiple merger coalescent models however, this assumption is less acceptable since the multiple merger events can in general affect long-range dependence within chromosomes (see Koskela [2018], Korfmann et al. [2023]). By computing the weighted SFS from this collection of SNP matrices, we can finely estimate rates of multiple merger events in the Λ-coalescent.s

## Summary of the method

In order to obtain a non-parametric estimator of the characteristic measure Λ, one has to follow these steps, that will be explained in detail in the next section.

1. Using a subsampling technique, that is explained in section 2.3, compute the weighted SFS for each of the *n* SNP matrices.
2. By averaging over the weighted SFS obtained from the different SNP matrices, obtain an estimator of the branch lengths of the tree, as given in (10).
3. Solving the linear system in Proposition 2.1, obtain estimates of the rates of multiple merger events in the Λ-coalescent.
4. Use a kernel based reconstruction (see (7)) to estimate the density of the characterizing measure of the coalescent. Consistency of this estimator is proven in Theorem 2.1.

## 2 Results

Table 1 summarizes the notations introduced in this section and used through the paper.

**Table 1:** Main notations

|  |  |
| --- | --- |
| <u>Integers/ indices:</u> |  |
| $m$ | number of individuals (or haplotypes) in the sample |
| $n$ | number of (almost) independent observations |
| <u>Multiple merger coalescents:</u> |  |
| $\Lambda, \nu$ | characteristic measure ($\Lambda(dx) = x^2 \nu(dx)$) |
| $\lambda_{k,j}$ | rates of multiple merger events (or coalescent rates) |
| $r_{k,j}$ | combinatorial coalescent rates |
| $d_k$ | sum of the combinatorial coalescent rates |
| <u>Density reconstruction:</u> |  |
| $(B_j^m)_{0 \leq j \leq m}$ | Bernstein polynomials of degree $m$ |
| $h$ | window parameter |
| $\phi$ | regularizing function |
| $K_h$ | kernel of window $h$ |
| $K_h^x$ | kernel of window $h$ around $x$ |
| $P_{m,h}^x$ | sequence of Bernstein polynomials used to approximate $K_h^x$ |
| $\nu_{m,h}$ | estimator of the density $\nu$ |
| <u>Estimator of the coalescent rates:</u> |  |
| $\hat{r}_{k,j}$ | estimator of the combinatorial coalescent rates |
| $\bar{c}$ | any partition of the integer $m$ |
| $\bar{c}_0$ | partition made of singletons (denotes the usual initial configuration of the coalescent process) |
| $L_j(\bar{c}_0)$ | partial length of order $j$ |
| $X_j$ | genotype of individual $j$ |
| $X_{j,k}$ | allele carried by individual $j$ at locus $k$ (0: wild-type, 1: mutated) |
| $(M_1(\bar{c}_0), \dots, M_{m-1}(\bar{c}_0))$ | site frequency spectrum of a sample of size $m$ |
| $(M_1(\bar{c}), \dots, M_{m-1}(\bar{c}))$ | weighted site frequency spectrum with weights $\bar{c}$ |

**Table 2:** Example of a SNP matrix, where 0 denotes the ancestral type and 1 denotes the derived type. Here the sample size is *m* = 6 and the total number of mutations is 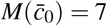.

| Ind. | SNP 1 | SNP 2 | SNP 3 | SNP 4 | SNP 5 | SNP 6 | SNP 7 |
| --- | --- | --- | --- | --- | --- | --- | --- |
| $X_1$ | 1 | 0 | 0 | 0 | 0 | 0 | 1 |
| $X_2$ | 1 | 0 | 0 | 0 | 0 | 1 | 1 |
| $X_3$ | 1 | 0 | 0 | 0 | 0 | 1 | 1 |
| $X_4$ | 0 | 0 | 1 | 1 | 0 | 0 | 0 |
| $X_5$ | 0 | 0 | 1 | 1 | 1 | 0 | 0 |
| $X_6$ | 0 | 1 | 0 | 0 | 0 | 0 | 0 |

### 2.1 Definition of the estimator

The Λ-coalescents are exchangeable continuous-time Markov chains taking their values in the partitions of the integers. The chain jumps from one state to another by coagulating some blocks of the partition to form one new block. The exchangeability implies that the jump rates only depend on the number of blocks of the partitions. Thus, the dynamics of this chain are totally described by the array (*λ*_*k, j*_)_*k≥ j≥*2_ where *λ*_*k, j*_ stands for the rate at which a given *j*-tuple, among *k* present blocks, coalesces into one new block. Moreover, the class of exchangeable coalescents is consistent. This means that a coalescent starting from a partition *π* made of *n* blocks, and restricted to the set of partitions of [*m*], with *m ≤ n*, has the same law as a coalescent starting from the partition *π* restricted to [*m*]. This consistency property implies that, for *k ≥ j ≥* 2,

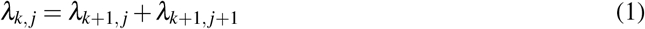

which is equivalent to

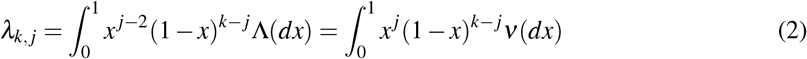

where *ν* is a measure on [0, 1] satisfying 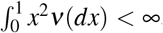, and Λ(*dx*) = *x*^2^*ν*(*dx*) (see Pitman [1999]). The measure Λ (or *ν*) fully describes the coalescent, in the sense that all the multiple merger rates can be computed by integrating against these measures, as shown by equation (2). One has to note that both measures Λ and *ν* characterize the coalescent, nevertheless the measure *ν* is easier to estimate than the measure Λ. Besides this fact, the estimation of *ν* can only insure a good estimation outside of an open interval containing 0 due to the presence of *x*^*−*2^ in its definition. In this work, we suppose that the measure *ν* is absolutely continuous with respect to the Lebesgue measure over the interval (0, 1], that is *ν*(*dx*) = *ν*(*x*)*dx*. This assumption includes Beta coalescents, but excludes Dirac coalescents as well as the Kingman coalescent.

The rate at which the number of blocks of the coalescent jumps from *k* to *j* is

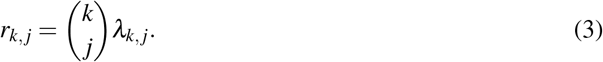

These new rates are then referred to as the combinatorial coalescent rates and the total coalescent rate from a state with *k* blocks is given by

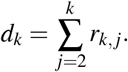

This section is focused on the practical reconstruction of the measure *ν* when estimations 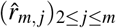 of the (*r*_*m, j*_)_2*≤ j≤m*_ are given. Note that the number *m* is fixed so that the only information needed is a good estimation of the coalescent rates of a group of *m* individuals. The reconstruction of the density of the measure *ν* depends on a hyperparameter *h* that can be understood as a parameter controlling the smoothing. Mathematically, this reconstruction can be seen as a map 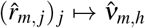 taking values in the continuous functions on (0, 1], where *h* holds for a window hyperparameter in a kernel estimation technique that we explain in details below.

The particular form of the coefficient *λ*_*m, j*_ obtained from (2) allows us to use Bernstein polynomials as a technical tool for the approximation step. The Bernstein polynomial basis is defined as a double index sequence given by

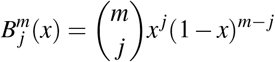

for all 0 *≤ j ≤ m*. In the following, we call Bernstein polynomial of order *m* a polynomial that belongs to the vector space spanned by the 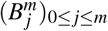 where *m* is fixed. The ideas behind the construction of the estimator 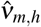 of *ν* are the following.

1. Choose a non-negative even function *φ* that is continuous, has a compact support containing 0 and such that ∫_R_ *φ* (*x*)*dx* = 1. Such a function is called a regularizing function. Without loss of generality, we assume that the support of *φ* is included in a segment of the form [*−τ, τ*]. For *h >* 0, define *K*_*h*_(*x*) = *h*^*−*1^*φ* (*h*^*−*1^*x*). The function *K*_*h*_ is called a kernel of window *h*. Classical functional theorems show that a continuous function *f* is well approximated by its smoothed version given by the convolution *f * K*_*h*_ defined, for any fixed *x*, by

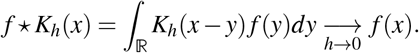 The convolution will apply on the function *ν* that has support included in (0, 1] so the integration of the function 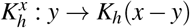 is only on the segment (0, 1]. The parameter *h* is a smoothing parameter and it has to be chosen carefully to avoid underfitting (*h* too small) or overfitting (*h* too large). Then, the kernel 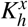 is a function that selects - in a smooth way - data points around *x*. The kernel itself is used to give weight to those points depending on their distance to the center point *x*. In general, a gaussian kernel (see section 3) is usually a good choice for pratical reasons but many more kernels can be used in this step. In the formal theorem given below (see Theorem 2.1), the bounded support condition is important for technical reasons but the gaussian kernel mimics this assumption and then works in practice. One can see [Györfi et al., chapter 5] for a detailed discussion on the consistency of kernel estimators.
2. Since the function 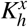 is smooth on its compact support, it can be approximated by a sequence of Bernstein polynomials 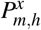 so that

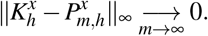 Those polynomials have the explicit definition

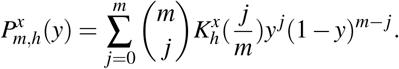
3. According to points 1. and 2., the quantity

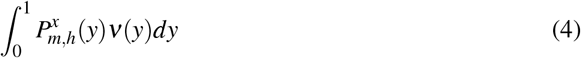

is a good candidate for the estimator of the density of *ν*(*x*). From the definition in (4), it is not clear to see that it is completely defined by the values of the *r*_*m, j*_. But it can be rewritten in the following way

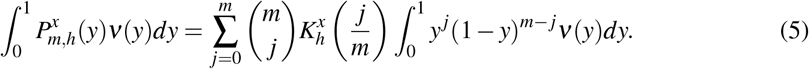 In the sum of the right hand side, the terms for *j* = 0 or *j* = 1 won’t lead to quantities that only depend on *r*_*m*_ = (*r*_*m, j*_)_2*≤ j≤m*_. This is not a problem if one takes *x* to be outside of the support of *K*_*h*_ for the terms *j* = 0, 1. This leads to the restriction *x−m*^*−*1^ *≥ hτ*. Then, we define

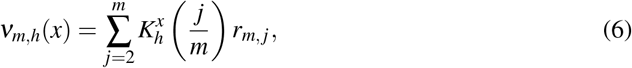

if *x ∈* [*m*^*−*1^ + *τh*, 1] and 0 elsewhere.
4. One has to note that *ν*_*m,h*_(*x*) is still not an estimator of *ν*(*x*) since it is not computable from the data. In the next section, we explain how to compute estimators of the coalescent rates 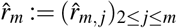. From (5) and (4), for any *x ∈* [*m*^*−*1^ + *τh*, 1], we define the final estimator 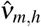 by a plug in of the estimated values 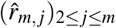,

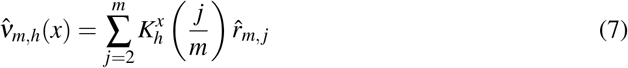

and we define 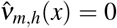 for *x < m*^*−*1^ + *τh*. In particular, we see that the interval [*m*^*−*1^ + *τh*, 1] tends to (0, 1] when we let *m →* ∞ and *h →* 0.

We establish in Theorem 2.1 that this estimator is consistent and we provide some control on the error. For that purpose we need to define the modulus of continuity. For a function *f* : [0, 1] *→* ℝ we define, for any *δ >* 0,

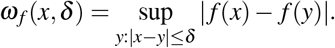

#### Theorem 2.1.

*Let τ >* 0, *h >* 0 *and m ≥* 2. *Let φ be a non-negative α-Hölder function of support* [*−τ, τ*] *and such that φ* (*x*) *≤* 𝟙_*{*[*−τ,τ*]*}*_(*x*). *Define K*_*h*_(*x*) = *h*^*−*1^*φ* (*h*^*−*1^*x*) *and for x ∈* [*m*^*−*1^ + *τh*, 1], *let* 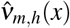 *as in* (7). *Then*,

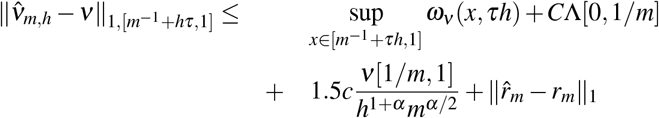

*where* 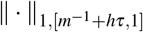 *holds for the L*_1_ *norm on the interval* [*m*^*−*1^ + *hτ*, 1], *and C, c are some positive constants*.

The proof of this Theorem can be found in Appendix A.

In words, there are four different sources of error in the reconstruction of *ν*. The first one comes from the modulus of continuity of *ν*. Since the measure *ν* is continuous, this quantifies how much it varies in a small interval of size *τh*. This is the most general way of stating the theorem. In most of practical cases, the function *ν* is sufficiently regular to weaken this assumption. In particular, if *ν* is Lipschitz of constant *k*, we can replace the term *ω*_*ν*_ (*x, δ*) by *kδ* when *x* and *δ* are such that *x − δ ≥ m*^*−*1^ + *τh* to insure to avoid any problem of discontinuity. Taking *δ* = *τh*, the window parameter *h* can be seen as a smoothing effect parameter. As every smoothing parameter it will have to be balanced in such a way to avoid over-estimation or under-estimation. This first term in the bound drives us in taking *h* the smallest possible whereas the third term in the bound is governing *h* the other way around. The second source of error comes from the mass of Λ close to 0. The third one to the mass of *ν* in [1*/m*, 1]. Observe that, since 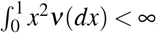, we have that *ν*[1*/m*, 1] = *o*(*m*^2^). This implies that we can always find a Hölder exponent *α* such that the term *ν*[1*/m*, 1]*/*(*h*^1+*α*^ *m*^*α/*2^) converges to 0. This also reflects the trade-off between the regularity of *φ* and the behavior of *ν* near 0. The fourth source of error is error in the estimation of the *r*_*m, j*_’s. The next section is devoted to providing a method to estimate these coalescent rates.

### 2.2 Techniques for the estimation of the coalescent rates

Let us turn to the problem of estimating the *m −* 1 combinatorial coalescent rates (*r*_*m, j*_)_2*≤ j≤m*_. The estimator 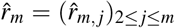 is defined as the solution of a linear system of *m −* 1 equations. Set *e* _*j*_ = (0,…, 0, 1, 0, …, 0) where the value 1 is at *j*-th coordinate.

To do so, we consider the set of partitions of the integer *m*, 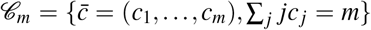. We denote the number of blocks of 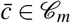 by 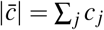. Then, we define the multi-dimensional block counting process of the coalescent with initial configuration 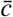 as 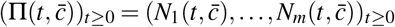, where 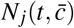 holds for the number of blocks of size *j* in the coalescent at time *t*. The total number of blocks in the coalescent process at time *t* is then given by 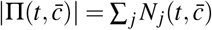.

The partial length of order *j* (1 *≤ j ≤ m−* 1) of the coalescent starting from 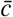 is defined by

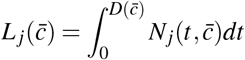

where 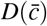 is the depth of the tree, also called time to the most recent common ancestor (TMRCA), 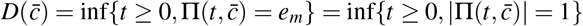.

#### Proposition 2.1.

*Let* 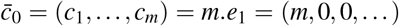 *and* 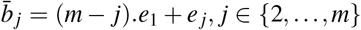. *Then we have. for i ∈ {*2,…, *m}*,

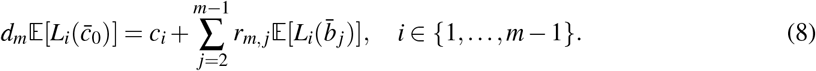

Equations (8) provide a linear system of *m−* 1 equations for *m−* 1 unknown values (*r*_*m, j*_)_2*≤ j≤m*_ which can be rewritten as

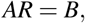

where *A* is a (*m −* 1) *×* (*m −* 1) matrix such that the last column contains the expected branch lengths, 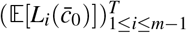 and for *j ∈ {*1, *m−* 2*}*, the *j*-th column contains the difference between the expected branch lengths and partial lengths starting from *b* _*j*_, i.e. 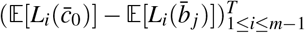; *B* is the vector (*m*, 0, …, 0) and *R* = (*r*_*m, j*_)_2*≤ j≤m*_.

The precision in the estimation of the vector *r*_*m*_ will thus depend on the precision of the estimation of the expectations in (8). The proof of Proposition 2.1 is a consequence of the Markov property applied at the first jump time (see Appendix B).

### 2.3 Obtaining the partial lengths from the SNP matrix

In this section we provide a method to estimate the expected values in (8). We assume that mutations happen at rate *θ*, as a Poisson process along the branches of the genealogical tree resulting from the coalescent process considered in the previous section. ^2^ This mutation rate can also be seen as a scaling parameter since the law of the mutations is not changed if one multiplies this parameter *θ* and the multiple merger rates are all multiplied by a common constant. Since the data is only considering the mutations affecting individuals in the sample, this scaling effect is untraceable, resulting in the model being not identifiable. To avoid dealing with those issues, we fix *θ* to be 1 in the sequel, which corresponds to measuring time in units of mutation rate. In the infinite sites model, we suppose that each mutation affects a distinct site so that they can all be observed in present-time data. This is a reasonable assumption since in most species the mutation rate is low, i.e. mutations are in nature rare events. In addition, multiple mergers generally reduce the time to the most recent common ancestor, which reduces the number of generations at which mutation events can occur.

We assume provided a sample of *m* individuals, whose genotypes are encoded in vectors *X*_1_, …, *X*_*m*_. Formally, the vector *X*_*j*_ describing the genotype of individual *j* is a vector *X*_*j*_ = (*X*_*j,k*_)_*k*_ *∈ {*0, 1*}*^*p*^, *p ∈* ℕ of binary values where *X*_*j,k*_ = 1 if and only if the individual *j* carries the mutation at site *k* (also called the SNP *k*). All this information is stored in a SNP matrix, where the vectors *X*_*j*_ are written as rows. See Table 2 for an example. Observe that the size of the vectors is not relevant here as the matrix can be adjusted by withdrawing columns of 0’s, which are the sites where no genetic variation is detected.

We also define the site frequency spectrum 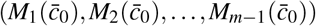 as the vector such that 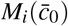 counts the number of mutations carried by exactly *i* individuals, that is 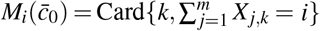. Also, the total number of mutations observed in the sample is 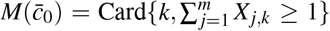. Notice that these quantities depend on the initial configuration 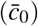. In fact, considering a sample of *m* different individuals corresponds to starting the coalescent process from *m* singletons, i.e. from 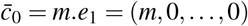. Our assumption on *θ* = 1 leads to

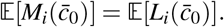

As a result, any estimator of the expected value 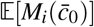 is also an estimator of the expected value of the partial length of order *i*.

In order to make use of the linear system (8), one has to be able to estimate the partial length 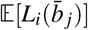 of a coalescent tree started at any partition 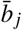 of the integer *m* of the form 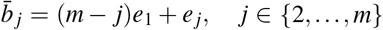. This can be done by a trick that consists in observing the site frequency spectrum differently and that is based on the consistency and the Markov properties of the coalescent. Indeed, the law of the coalescent tree starting from a partition 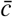 is the same as that of the coalescent tree starting from 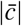 individuals where individuals are associated to weights given by the partition 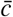. We will therefore define a *weighted* SFS, by subsampling *m*′ individuals with replacement. The weight of each individual is given by the number of times it appears in the subsample. A mutation carried by an individual with weight *j* will count as a mutation carried by *j* individuals. The weighted SFS corresponding to a subsample with weights 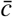 is denoted by 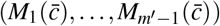.

Let us give an example using the SNP matrix of Table 2. In this matrix *m* = 6, 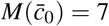 and the observed SFS is (2, 3, 2, 0, 0). Now, think of 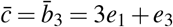, i.e. a sample where three individuals have weight 1 and one individual has weight 3. We chose at random one individual to which we associate the block of size 3, e.g. *X*_1_ and three individuals to which we associate weight 1 (*X*_2_, *X*_3_ and *X*_4_). Then the weighted version of *X*_1_ is (3, 0, 0, 0, 0, 0, 3). The corresponding weighted SFS is (2, 1, 0, 0, 2). Recall that, due to the exchangeability of the coalescent process, the law of the weighted SFS does not depend on the labels of the individuals which are assigned to each weight.

Formally, for a general partition 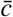 of any integer *m*′ *≥ m* such that 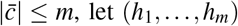, let (*h*_1_, …, *h*_*m*_) be any partition vector associated to 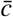, such that 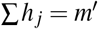, Card*{ j, h* _*j*_ = *i}* = *c*_*i*_ and 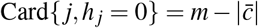. Then,

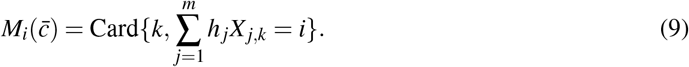

Observe that *m*′ = *m* in the example above.

Now, suppose that we can observe *n* i.i.d. SNP matrices of *m* individuals and that from each of these *n* matrices we obtain a realization of 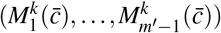 for *k ∈ {*1,…, *n}* using the subsampling procedure described above. The estimator of 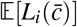 is given by

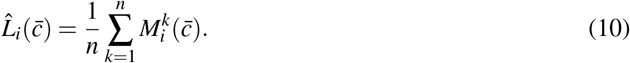

## 3 A numerical example

We apply our method to simulated data, where the coagulation measure Λ is known, so we can compare the estimated measure 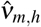 to the true measure *ν*. The goal of this section is not to quantify the error in the reconstruction, which is already bounded in Theorem 2.1, but rather to illustrate our method.

We used the population genetics simulator `msprime` [Baumdicker et al., 2022, Kelleher et al., 2016]. There are two Λ-coalescent models available in `msprime`: the Beta-coalescent [Schweinsberg, 2003] and the Dirac-coalescents [Eldon and Wakeley, 2006]. Since we have assumed that the Λ measure has a (Lebesgue) density, we chose the Beta-coalescent model. It is a one parameter family in which the characterizing parameter *α* denotes the degree of skewness of the offspring distribution of the population. Smaller values of *α* correspond to a more skewed offspring distribution, while *α →* 2 corresponds to the Kingman coalescent. For this example we fixed *α* = 1.2, but we obtain qualitatively similar results for other values such that 1 *< α <* 2, which is the range of values allowed by `msprime` (Supplementary Figures S2 and S3). Observe that in this model the characteristic measure *ν*(*x*) behaves like *x*^*−*1*−α*^ near 0, which implies it has no first moment, providing a “worst-case scenario” for the non-parametric estimation.

We simulated the SNP matrix of a sample of size *m*. We estimated the partial lengths by computing the SFS and the weighted SFSs, as explained in the previous section. We simulated *n* independent SNP matrices, that we used for inference. We explored different settings for *m* and *n*. Then we solved the linear system (8) in Proposition 2.1. Since some of the coalescent rates (the *r*_*m, j*_’s) are small, sometimes solving the linear system *AR* = *B* with the estimated values of the partial lengths 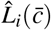’s yielded negative values of the *r*_*m, j*_’s, so, instead we used `scipy.optimize`, to minimize *AR − B* with the constraint that the *r*_*m, j*_’s must be positive.

To reconstruct the measure *ν*, we followed our method using a Gaussian kernel for *K*_*h*_. Figure 1 compares the true density of *ν* and the density of *ν* reconstructed using the true values of the *r*_*m, j*_’s (using equation (6)). We can observe that there is a rather large error close to 0. This is because the family of Λ measures that we were able to simulate using `msprime` have a lot of mass close to 0 (which corresponds to the second term of the error computed in Theorem 2.1). We show results for two different values of *h*. We observe that the error in the reconstruction depends on *h*, and that for *m* = 50, *h* = 0.02 is a good parameter choice.

**Figure 1:**
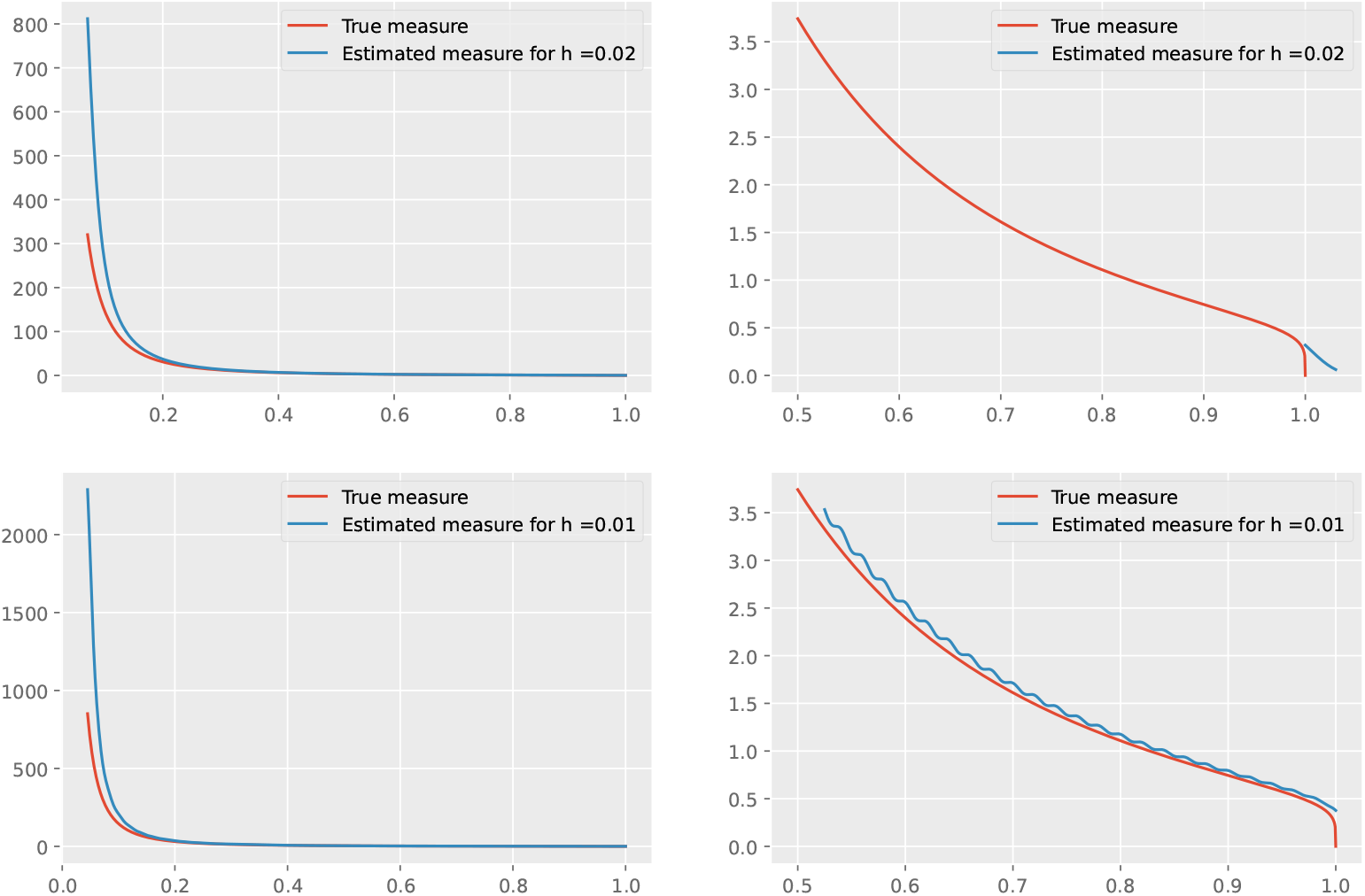
Density of *ν*. Here the true value of the coalescent rates (*r*_*m, j*_’s) is used to reconstruct the density of *ν* using equation (6) (blue curves). We compare against the true density of the measure *ν* (red curves). We used a a Beta-coalescent model with *α* = 1.2 and *m* = 50 and we show two different values of *h*. Panels on the right are zoomed versions of the panels on the left (notice the difference in the x-axis).

In a second step, we applied the whole method to our simulated data, i.e. we first estimated the *r*_*m, j*_’s from our simulated SNP matrices and then we reconstructed the measure *ν*.

We simulated different values of the sample size *m* and (population-size rescaled) mutation rate *θ*. Our method does not allow to estimate the mutation rate *θ*, but as discussed in the previous section, changing *θ* corresponds to changing the time-scale, i.e. multiplying the *r*_*m, j*_’s by some constant. Since the aim of this project is not to get this constant, in order to visualize the results, we normalized the measures by the integral of Λ in [*m*^*−*1^ + *hτ*, 1], which is an approximation for the total coalescence rate, i.e. the constant we need.

We fixed the number of independent replicates to *n* = 20. This would correspond, for a species with 20 chromosomes, to consider that each chromosome is an almost independent replicate. Figure 2 shows the obtained results, which strongly depend on the mutation rate. For *θ* larger than 1 (which corresponds more or less to 500 mutations per sample in our model), the estimation of *ν* is quite good, in the sense that it has a similar behavior to the true *ν*. However, for lower mutation rates, the estimated measure does not match with the real measure.

**Figure 2:**
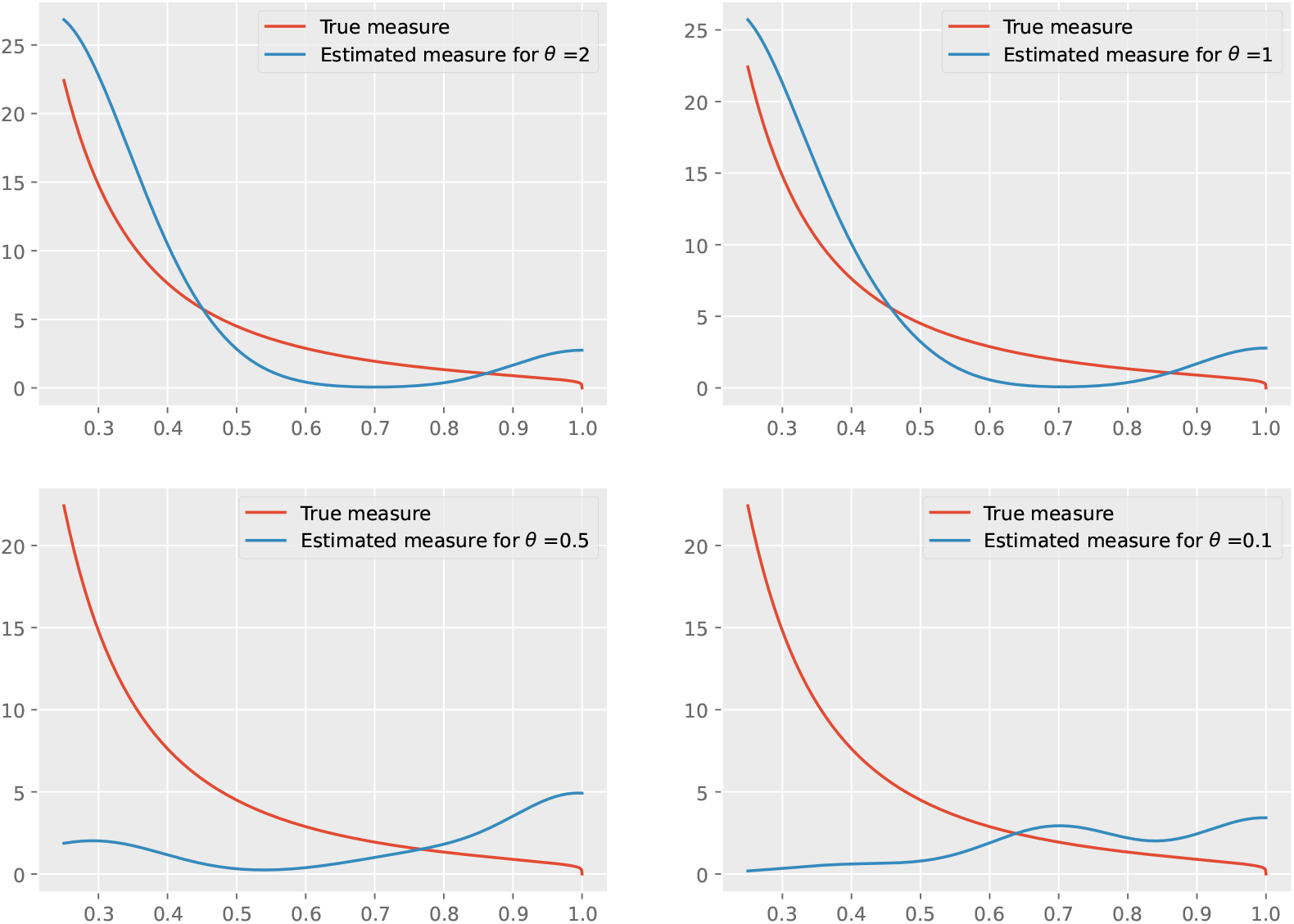
Density of *ν*. We compare the true density (red curves) to the density estimated from simulated SNP matrices using equation (7) (blue curves). Here *n* = 20, *m* = 10, *h* = 0.1, *τ* = 2, *α* = 1.2.

If we increase the number of independent samples, the quality of our estimates improves (Figure 3). However, obtaining such a large number of almost independent samples, with enough mutation on each requires either a high-mutation high-recombination regime that is difficult to observe from real data. This parameter regime seems more likely to be achieved if the independent samples are replicates from an evolutionary experiment.

**Figure 3:**
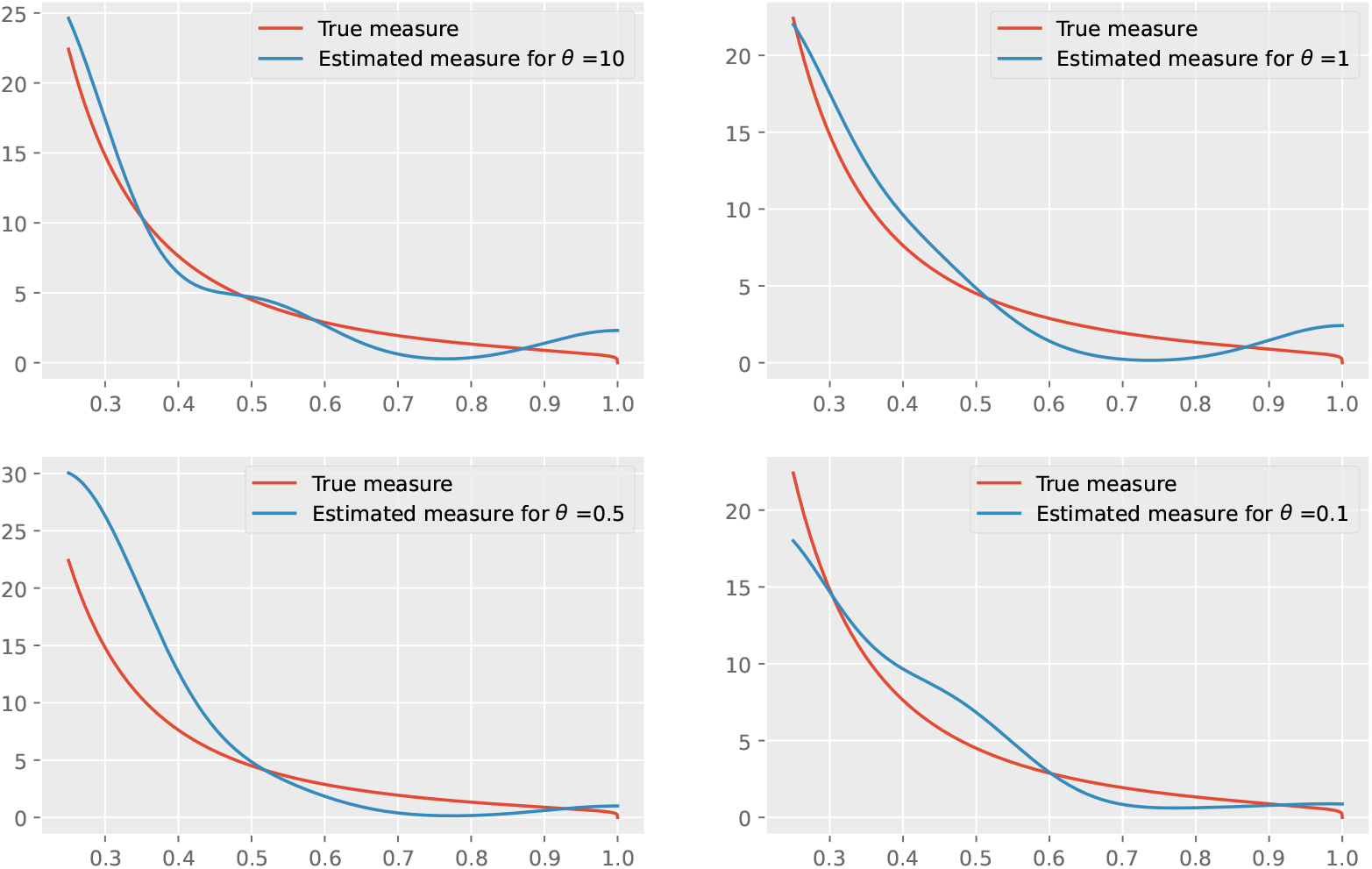
Density of *ν*. We compare the true density (red curves) to the density estimated from simulated SNP matrices using equation (7) (blue curves). Here *n* = 100, *m* = 10, *h* = 0.1, *τ* = 2, *α* = 1.2.

Supplementary Figure S1 shows the real values of the multiple merger rates *r*_*m, j*_’s and their estimated value 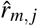’s. We can observe that, although the error in the estimated multiple merger rates is large (especially when *r* is close to *m*, the density of the reconstructed measure still shows the same trend as the true density.

Finally, increasing *m* does not necessarily provide better estimates (Supplementary Figures S6 and S7). This is because the effect of increasing the sample size is counteracted by an increase in the size of the linear system given by (8) and the errors in the estimation of the partial lengths propagate through the system. Decreasing *m* (Supplementary Figures S4 and S5) can also increase the reconstruction error (see Theorem 2.1).

## 4 Discussion

We have developed the first non-parametric estimator of the density function of the measure Λ of Λ-coalescents from observations taken at a single time point. Our method can infer any measure Λ, in a way that is agnostic to biology, since we are not making any assumption on how the population evolves. It could be used to hint or exclude already known coalescent classes such as Beta coalescents or the Bolthausen-Sznitman coalescent which are linked to biological phenomena such as skewed offspring distribution or certain modes of selection. It provides a well-fitting model for observed genetic diversity, which can be used to find species or genomic regions where other processes additionally, on top of those modelled by classical Λ-coalescents (Beta, Dirac or or the Bolthausen-Sznitman) influence genetic diversity. We are also able to quantify the consistency of this estimator in our main theorem. Note that our method fits better models with a measure Λ having a non-explosive behavior close to 0, providing a convincing rejection tool of the family of Beta coalescents.

The method works in three steps. First, we estimate the branch lengths of the coalescent from SNP matrices thanks to a new observable that we called the weighted SFS. Second, we obtain the coalescent rates from the branch lengths by inverting the linear system (8). Third, we estimate the coalescent measure from the coalescent rates thanks to a non-parametric method based on Bernstein polynomials, as explained in the first paragraph of the Results section. Our (new) mathematical results focus only on step 3. Notice that steps 1 and 2 may be useful for other statistical techniques, in particular if SFS based statistics are needed, or if coalescent rates are needed.

The error generated at step 1 does only come from the central limit theorem so it is minimized if we can observe data from a large number of independent replicates or if the mutation rate is high. It can increase a lot if those assumptions do not hold. We tried the weighted SFS method by by simulating genome-wide data from a single sample and treating distant genomic regions as independent (pseudo-

)samples. To do so, we first simulated the ancestry of a very large population using `msprime` and then subsampled individuals with or without replacement. This yielded large errors in the estimation of the expected branch lengths. This is because of the shared evolutionary history between individuals of a same population, that makes the ancestry trees of the different samples highly correlated (especially for the deeper nodes).

The error produced at step 2 is due to the propagation of the error in step 1 at the moment of inverting a linear system. This part might be improved by using some more efficient numerical tools, or by adding constraints or equations in the system, e.g. recursive equations derived from (1). A more promising track to follow is that we may need not all the rates *r*_*m, j*_ since they get too small when *j* goes close to *m* and do not bring substantial information for the estimation of Λ.

Finally, we managed to bound the error generated at step 3 in Theorem 2.1 thanks to the theory of non-parametric statistics. The error naturally explodes close to 0 when the measure Λ accumulates too much mass at 0. This is typically the case in Beta-coalescents with parameter *α* between 1 and 2. Thus, obtaining a non-explosive measure close to 0 could also mean that the evolution of the population does not fall into the domain of attraction of Beta coalescents.

In this work we did not focus on the estimation of the mutation rate *θ*. Recall that it is not possible to discriminate between the couple (*θ*, Λ) and (*cθ, c*Λ) for some constant *c* from one date set taken at a single time. We refer to Eldon et al. [2015], Freund and Siri-Jégousse [2021] for some further techniques involving estimation of the mutation rate and model selection.

## 5 Code availability

Code for the numerical example is available at : https://github.com/vmiropina/bootstrap_coalescent/.

## 7 Appendix A: Proof of Theorem 2.1

In this section we prove the consistency of the proposed estimator. One source of error comes from the uncertainty on the terms 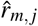. There are also two other sources of approximation error given by the fact that we approximate *ν* by *ν * K*_*h*_ and that we approximate 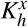 by 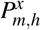. To prove the theorem, we need to tackle these three sources of error. The first results that we present deals with the smooth approximation of *ν* by *ν * K*_*h*_.

### Proposition 7.1.

*Let φ be such that for all x ∈* ℝ, *φ* (*x*) *≤* 𝟙_*{*[*−τ,τ*]*}*_(*x*), *for some τ >* 0. *Then, for any δ >* 0 *and any x ∈* (0, 1] *such that x−δ ≥ m*^*−*1^ + *τh*,

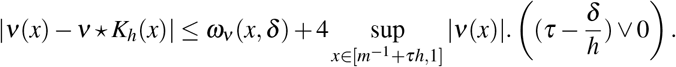

*Proof*. Using the fact that *K*_*h*_ is of integral 1, we see that for any *δ >* 0 and any *x*,

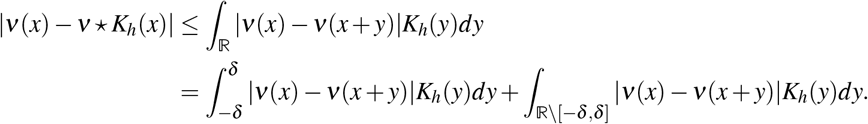

Observe that the last term is well defined since the support of *K*_*h*_ is included in [*−τh, τh*]. Then,

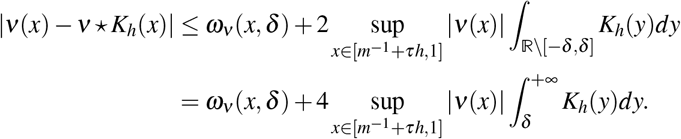

This last integral is obviously 0 if the support of *K*_*h*_ and [*δ*, +∞) are disjoint i.e., if *τh < δ*. Otherwise,

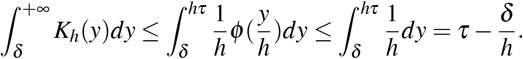

Next, we deal with the approximation of 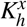 by the Bernstein polynomials 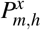.

### Proposition 7.2.

*Let f be a continuous function on* [0, 1]. *Then*

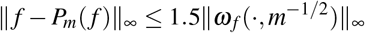

where 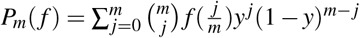

The proof can be found in Theorem II, p. 518 of Queffélec and Zuily [2020] where the constant can be taken equal to 1.5 from the proof page 519. Proposition 7.2 will be applied on the kernel function *K*_*h*_. It is obvious to see that for any *δ >* 0,

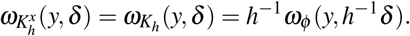

Thus the regularity of 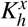 is directly inherited from the regularity of *φ*. For example, if *φ* is *α*-Hölder,

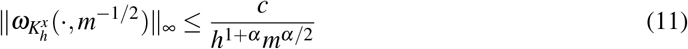

for some constant *c >* 0.

The following simple result is the verification that the plug in strategy of (7) is valid when one controls the *L*_1_ error of estimation on the vector *r*_*m*_ = (*r*_*m, j*_)_2*≤ j≤m*_.

### Proposition 7.3.

*For any h >* 0 *and any m ≥* 2 *one has that*

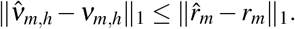

*Proof*. The proof of the proposition is obtained by noticing that the terms *K*_*h*_(*x− j/m*) have integral equal to 1. More formally, notice that the support of 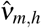 or *ν*_*m,h*_ is included in [*m*^*−*1^ + *τh*, 1] and so

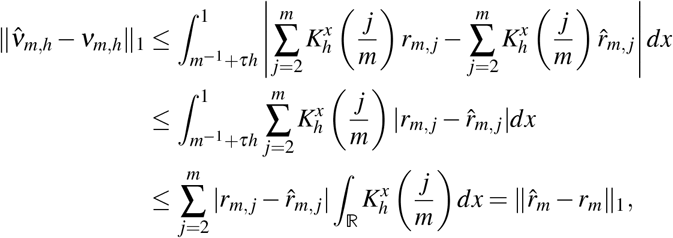

which concludes the proof.

When one sums up the previous results we end with the following result of consistency.

*Proof of Theorem 2.1*. Noticing that for any *x*,

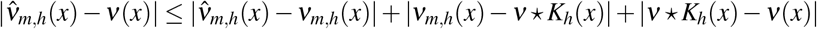

and using the previous results, we see that it is enough to show that

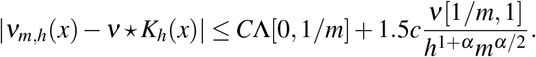

But since *x* belongs to [*m*^*−*1^ + *hτ*, 1], we have that 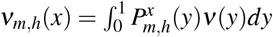 and so

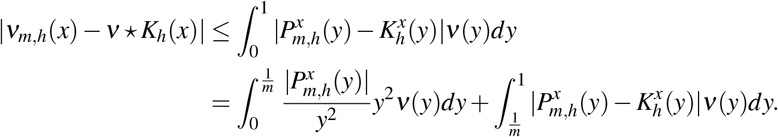

On the one hand, the polynomial 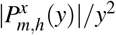 is uniformly bounded so there exists a constant *C* such that

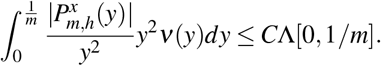

On the other hand, by Proposition 7.2 and (11), we have that

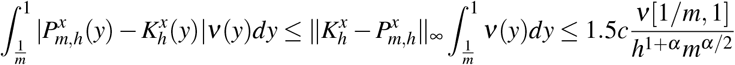

## 8 Appendix B: Proof of Proposition 2.1

In this section we prove Proposition 2.1 by establishing a more general recursion result. Consider any initial configuration 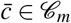. By the strong Markov property (applied at the time of the first jump of the process), we obtain that, for *i ∈ {*1,…, *m−* 1*}*,

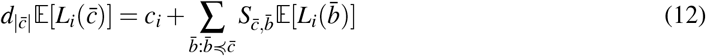

where 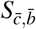 is the rate at which the multi-dimensional block counting process jumps from 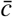 to 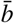, i.e.

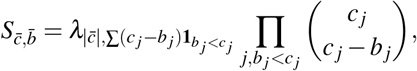

already defined in Eq. (20) of Hobolth et al. [2019]. In the summation in (12), the notation 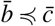 holds for the vectors 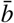 obtained by a single step merging of a multiple number of lineage into one. This imposes, in particular, that there exists a unique index *i*_0_ *≤ m* such that 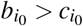 and in this case, 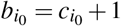. This last fact is actually a characterization of the partial order ≼. An additional fact is that

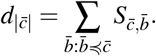

Proposition 2.1 is obtained in the particular example where the initial configuration only contains singletons, that is 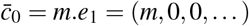. In this case we recover the classical functional leading to the site frequency spectrum. After the first merge, the *m−* 2 attainable partitions in (12) are of the form

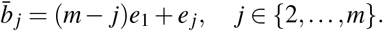

An important observation is that, in this case, 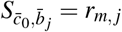.

### Remark 8.1.

*There is a classical relation between the rates* (*r*_*m, j*_)_2*≤ j≤m*_ *and the expectation of the total branch lengths of the associated coalescents starting from j ∈ {*2,…, *m} individuals, denoted by B*(*j*). *The total branch length is the sum of all the lengths of the coalescent tree from the leaves until the root. It is also obtained thanks to the Markov property applied at the first jump time:*

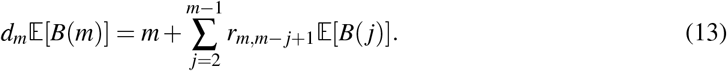

*A first approach to estimate the coalescence rates would consist in considering this equation and add a number of recursive equations* (1) *to obtain a linear system of m −* 1 *equations. However this very rigid strategy, based only on the total number of blocks of the coalescent, leads to uncontrolled error as the error resulting from the estimate of r*_*m*,2_ *will spread additively in the rest of the estimates. The strategy based on recursive equations for the partial lengths of the coalescent, as proposed in Proposition 2.1, leads to a more stable system*.

**Figure S1:**
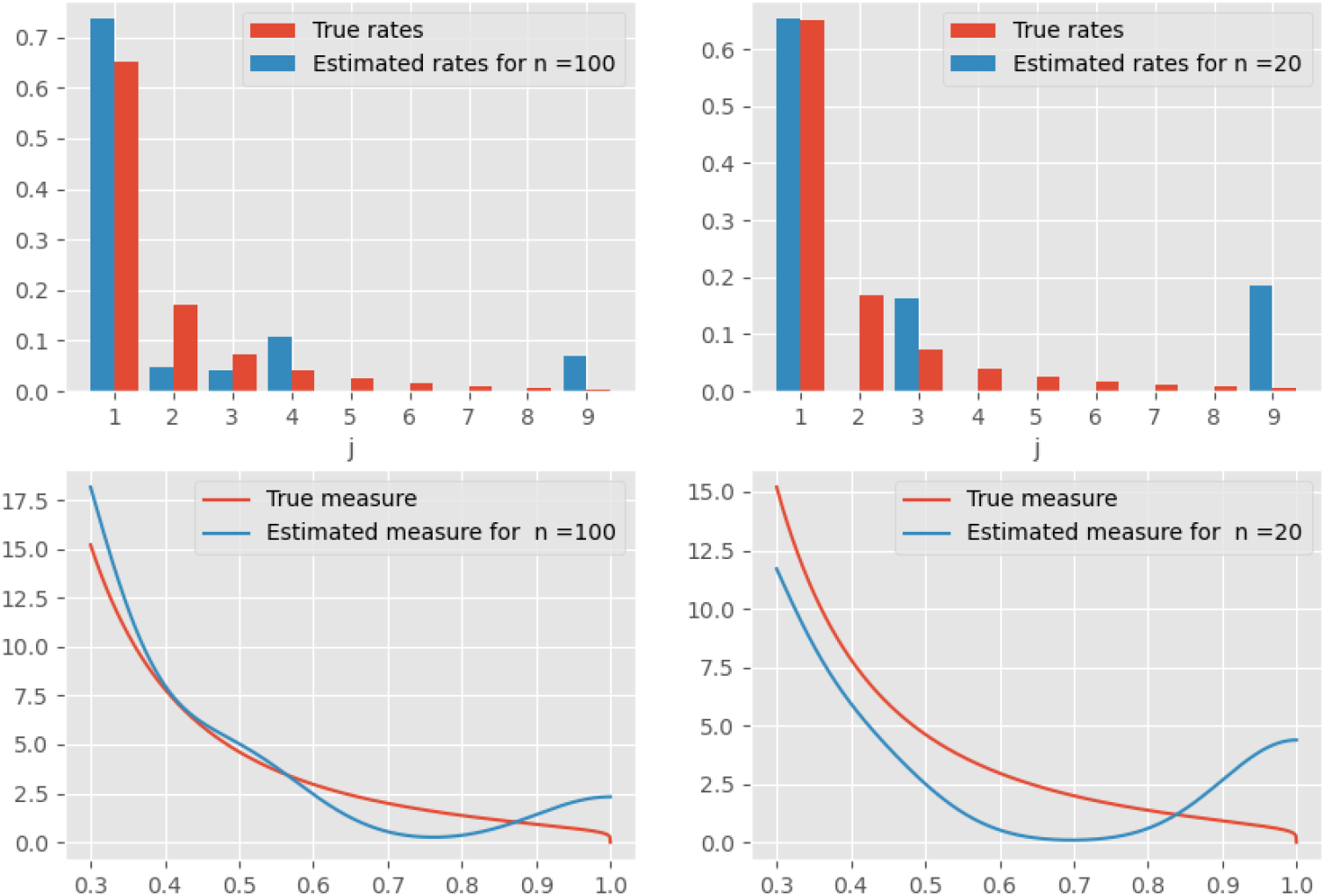
Multiple merger rates and density of *ν*. In the upper panels, we compare the multiple merger rates *r*_*m, j*_’s to the estimated values 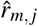’s In the bottom panels, we compare the true density (red curves) to the density estimated from simulated SNP matrices using equation (7) (blue curves). Here *n* = 20, *m* = 50, *h* = 0.01, *τ* = 2, *α* = 1.2.

## 9 Appendix C: Supplementary Figures

**Figure S2:**
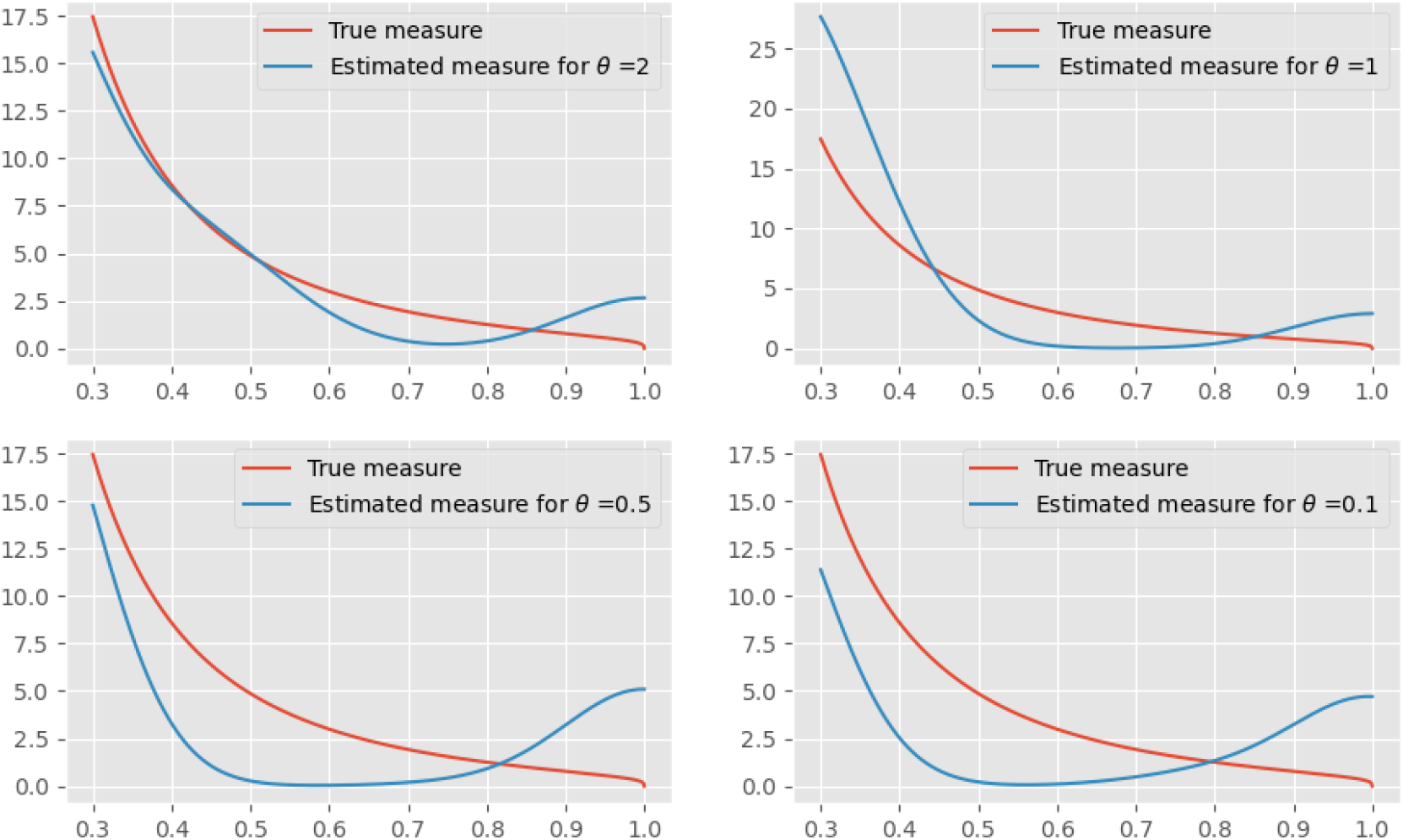
Density of *ν*. We compare the true density (red curves) to the density estimated from simulated SNP matrices using equation (7) (blue curves). Here *n* = 20, *m* = 50, *h* = 0.01, *τ* = 2, *α* = 1.3.

**Figure S3:**
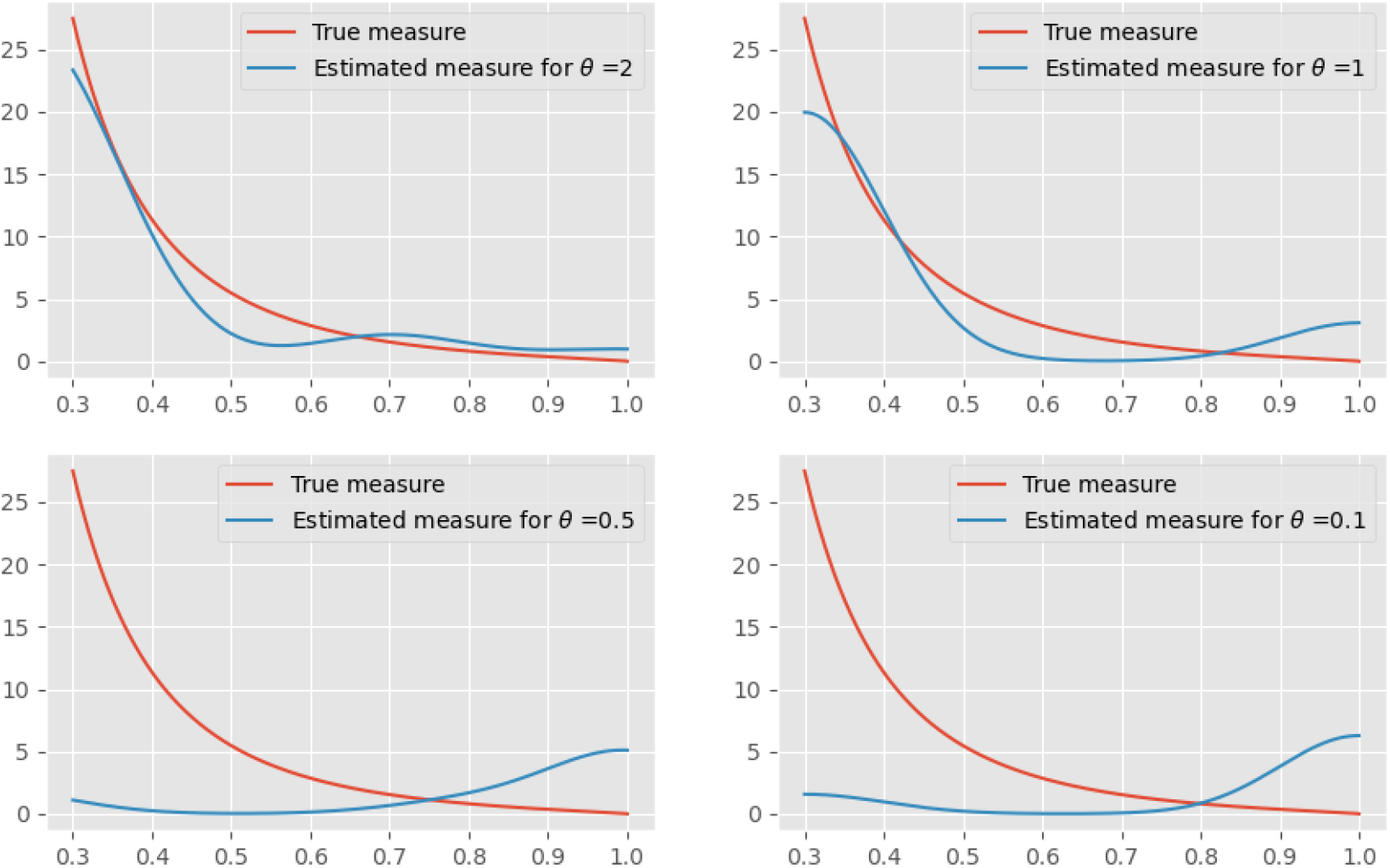
Density of *ν*. We compare the true density (red curves) to the density estimated from simulated SNP matrices using equation (7) (blue curves). Here *n* = 20, *m* = 50, *h* = 0.01, *τ* = 2, *α* = 1.7.

**Figure S4:**
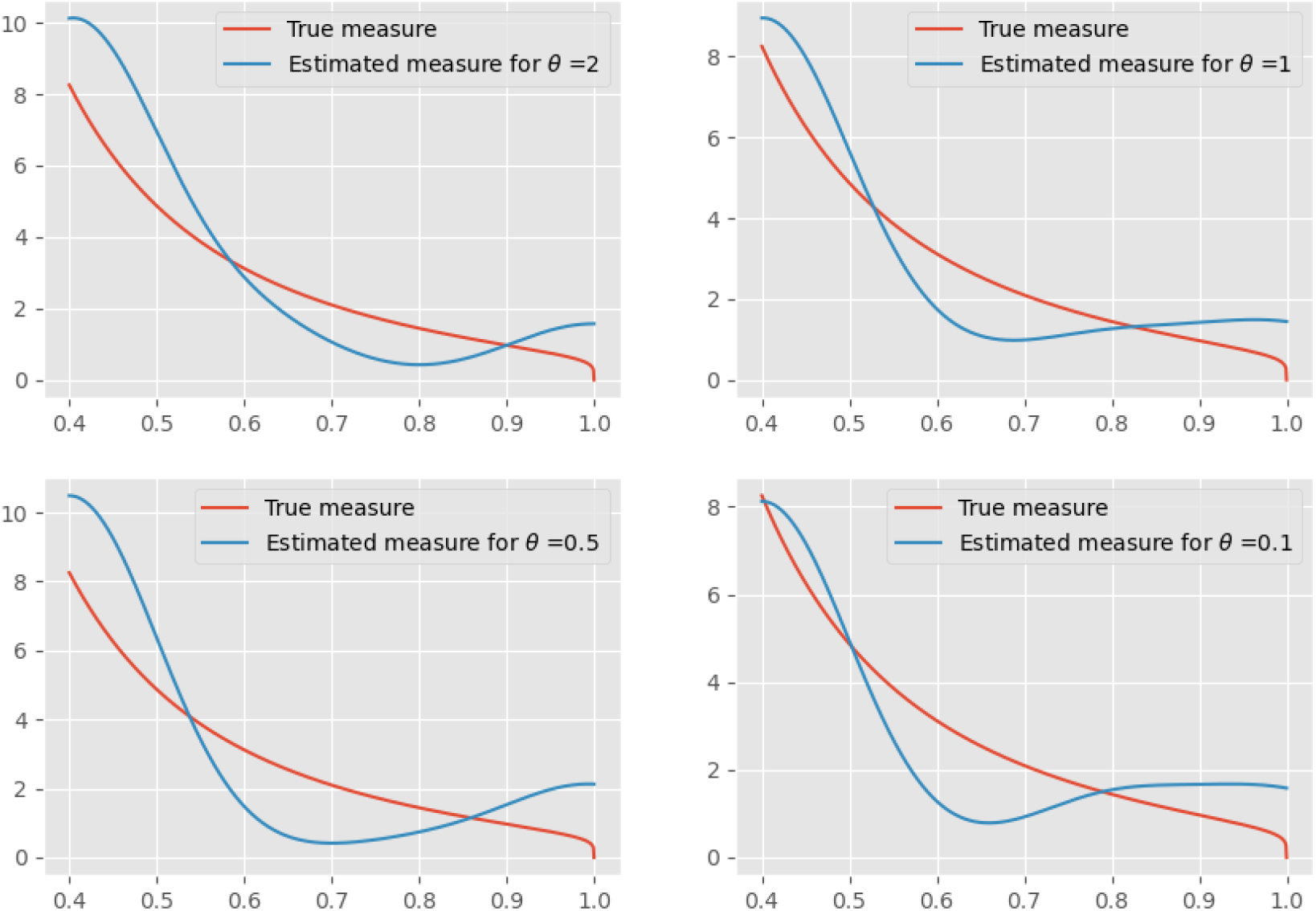
Density of *ν*. We compare the true density (red curves) to the density estimated from simulated SNP matrices using equation (7) (blue curves). Here *n* = 20, *m* = 5, *h* = 0.1, *τ* = 2, *α* = 1.2.

**Figure S5:**
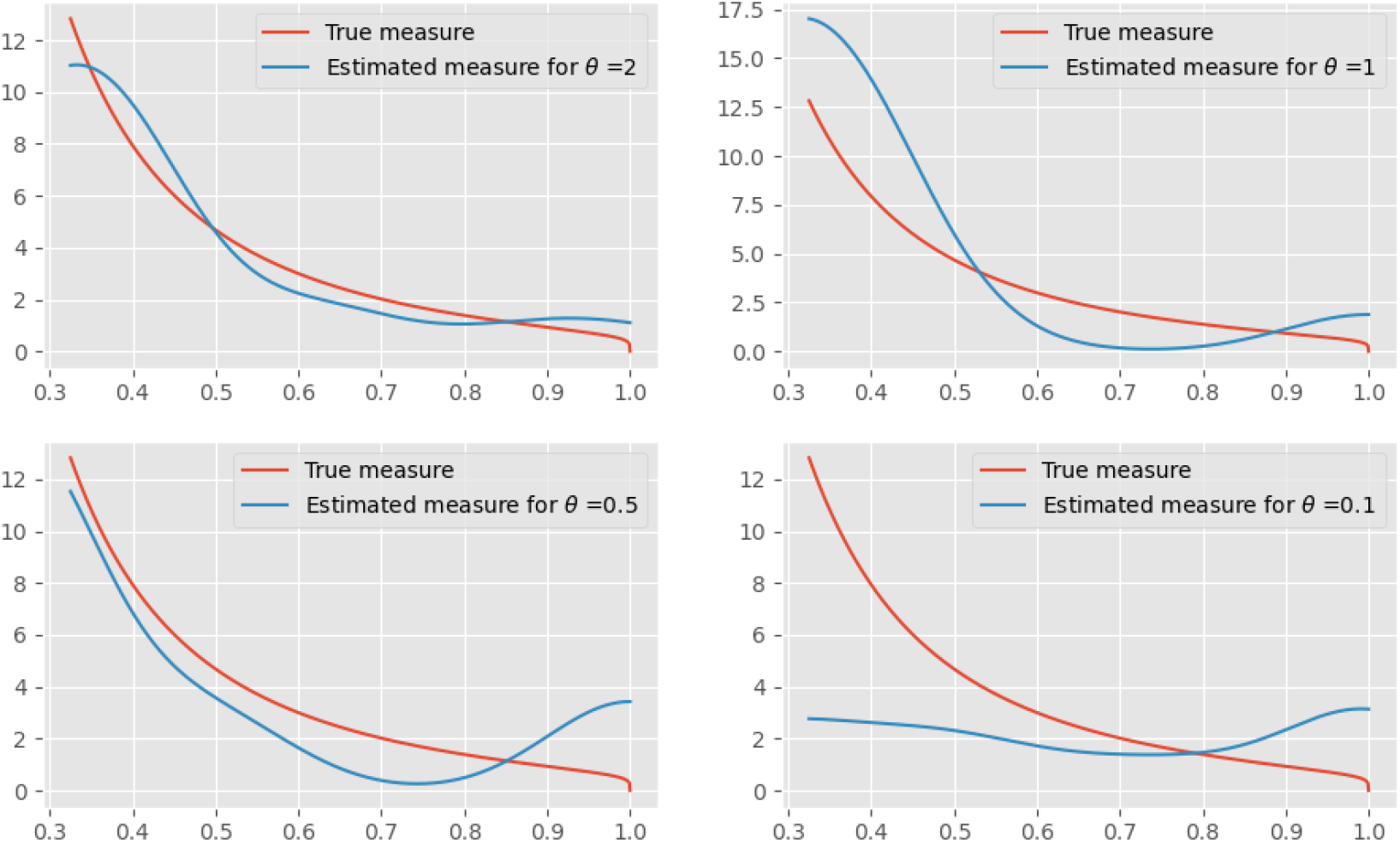
Density of *ν*. We compare the true density (red curves) to the density estimated from simulated SNP matrices using equation (7) (blue curves). Here *n* = 20, *m* = 8, *h* = 0.1, *τ* = 2, *α* = 1.2.

**Figure S6:**
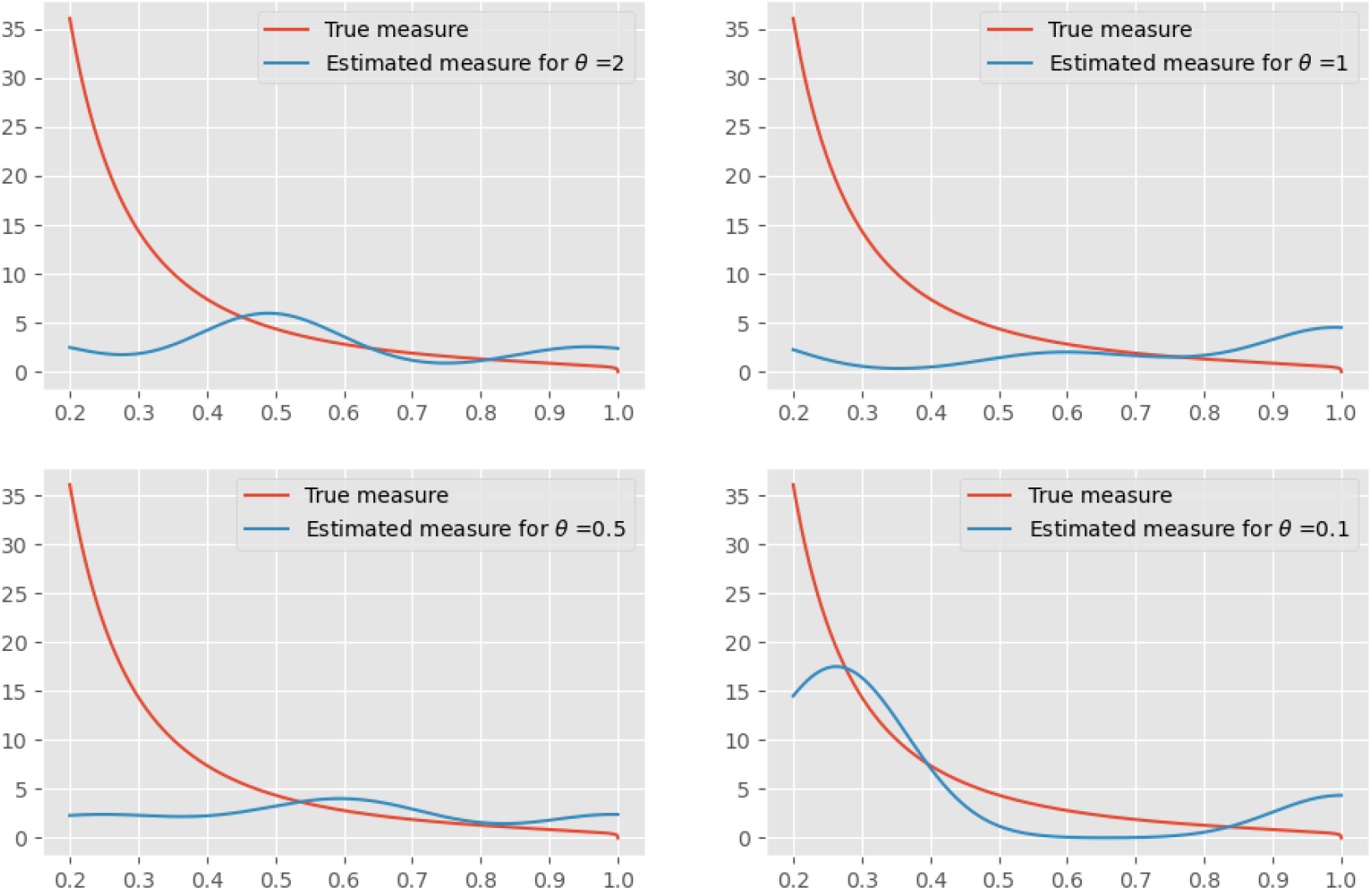
Density of *ν*. We compare the true density (red curves) to the density estimated from simulated SNP matrices using equation (7) (blue curves). Here *n* = 20, *m* = 20, *h* = 0.1, *τ* = 2, *α* = 1.2.

**Figure S7:**
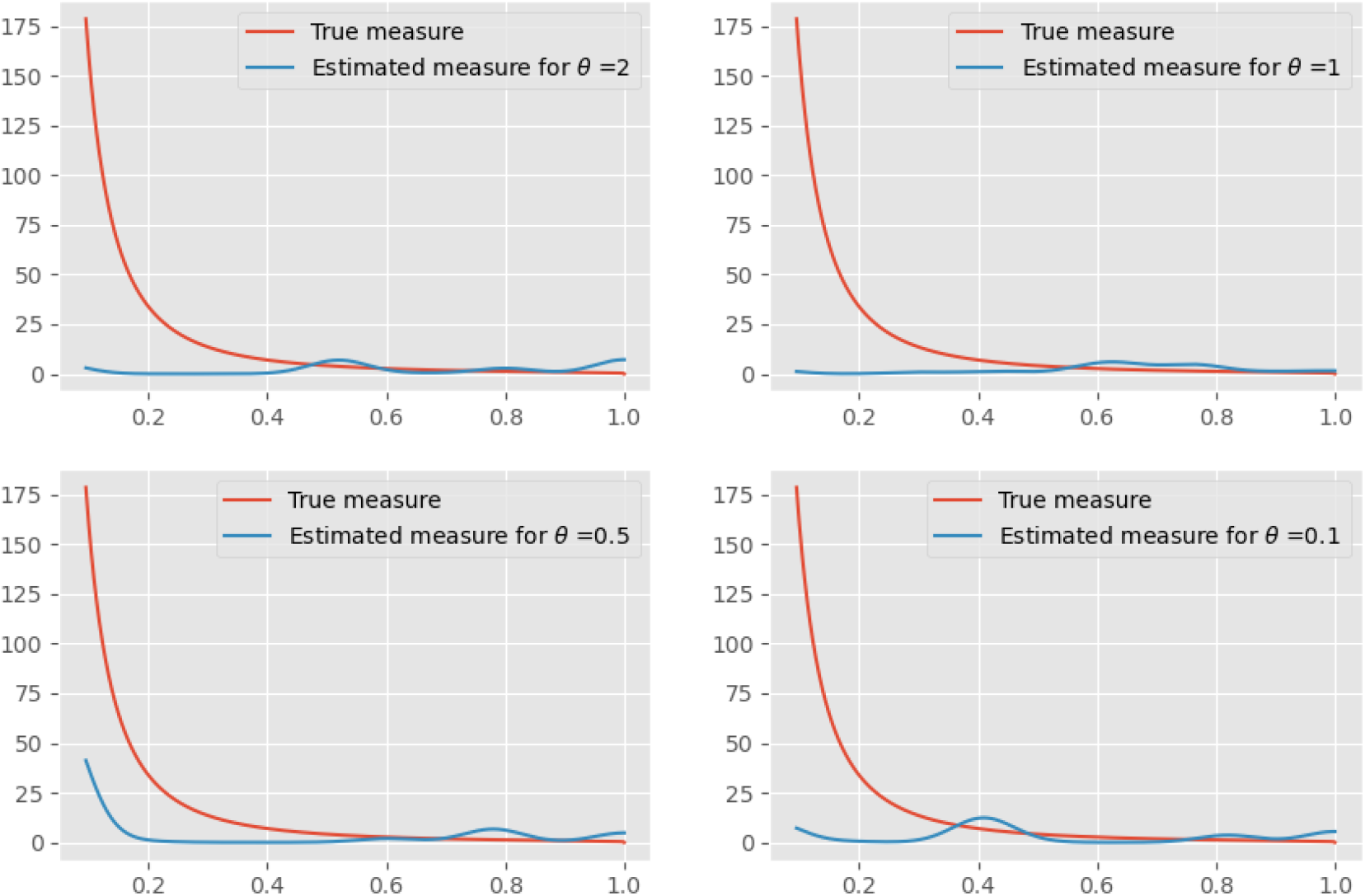
Density of *ν*. We compare the true density (red curves) to the density estimated from simulated SNP matrices using equation (7) (blue curves). Here *n* = 20, *m* = 50, *h* = 0.02, *τ* = 2, *α* = 1.2.

## Footnotes

1 Single Nucleotide Polymorphisms (SNP) are mutations that affect one single base position in the DNA. For the purpose of this paper, SNPs can be substitutions or indels.

2 *θ* is the population size rescaled mutation rate, i.e. *θ* is proportional to *N*_*e*_ *× μ*, where *N*_*e*_ is the effective population size and *μ* is the mutation rate of the sequence, per generation. In humans, the mutation rate per nucleotide per generation is of the order of 10^*−*8^ and *N*_*e*_ is of the order of 10^4^, which would correspond to *θ* = 1 if the sequence length is of order 10^4^.

